# Starting strong: A descending analgesic dosing scheme enhances pain relief compared to a conventional ascending dosing scheme

**DOI:** 10.64898/2026.10.10.758177

**Authors:** Marilyn Mintah, Jean Daunizeau, Alexandra Tinnermann

**Affiliations:** Institute of Systems Neuroscience, University Medical Center Hamburg-Eppendorf, Germany; Brain and Spine Institute, Paris, France

## Abstract

Managing pain while minimizing opioid-related harm remains a central clinical challenge. Enhancing drug-related placebo effects has been proposed as one strategy for reducing opioid consumption, yet current dosing guidelines which recommend starting analgesic treatment at a low dose and titrating upward may inadvertently generate negative treatment expectations, since initial drug encounters are often associated with minimal pain relief. This study tested whether the inverse approach of a descending dosing scheme that starts with a strong analgesic dose more effectively enhances placebo-related analgesia than the conventional ascending scheme. Eighty-two healthy participants underwent a heat pain paradigm involving a sham analgesic treatment delivered at three different efficacy levels (low, medium, high). Participants were randomly assigned to a descending (high-to-low) or ascending (low-to-high) treatment-history sequence before receiving an identical, medium-intensity treatment during a subsequent test phase. The descending group showed significantly greater pain relief during the test phase than the ascending group, despite receiving identical treatment. These findings suggest that initiating analgesic treatment with a high, effective dose can amplify placebo-related analgesia relative to conventional dosing schemes, offering a potential strategy for reducing cumulative opioid use without compromising pain relief.

## Introduction

Pain treatment is an important but also challenging endeavour. While one important goal is relieving patients from their pain, drug dependencies and addiction are a major risk when treating patients with analgesics such as opioids. Opioid addiction and opioid-related deaths have risen dramatically in recent years, driven largely by increased opioid prescriptions (Guy, 2017; Han et al., 2015; Ladha et al., 2019; Scholl, 2019). While most pronounced in the US, similar observations have also been reported in the UK, France, Australia, and Germany over the past two decades (Alenezi et al., 2021; Chenaf et al., 2019; Karanges et al., 2016; Rosner et al., 2019). These observations underscore the need for approaches that can lower opioid consumption while maintaining adequate pain control. Enhancing drug-related placebo effects represents one such approach for reducing the amount of administered opioids (Pattullo and Colloca, 2019). Numerous studies have shown that drug effects extend beyond what can be attributed to the chemical agent alone (Wager and Atlas, 2015). For example, when patients or participants are unaware that they are receiving an analgesic, pain relief is substantially reduced compared to when they know they are receiving an analgesic (Amanzio et al., 2001; Bingel et al., 2011; Kam-Hansen et al., 2014; Levine et al., 1981). These findings suggest that part of the drug-induced analgesia is driven by placebo effects that arise from positive treatment expectations and learning (Voudouris et al., 1990). Another example is pharmacological conditioning, where placebo effects can prolong pharmacological effects. Once analgesic treatments are covertly substituted with saline, analgesia in humans (Amanzio and Benedetti, 1999; Benedetti et al., 2007, 2003) and animals, including rats (Boorman and Keay, 2021a; Miller et al., 1990; Neyama et al., 2025) continues to persist. Notably, changing the dose during pharmacological conditioning has been shown to attenuate conditioned analgesia in animals (Boorman and Keay, 2021b), suggesting that dosing schemes themselves influence placebo-related expectations and learning and could be optimized to maximize drug-related placebo effects. The World Health Organization's “Pain Relief Ladder” for cancer pain treatment represents one such dosing scheme (World Health Organization, 1996) that recommends a stepwise escalation from non-opioid drugs to mild opioids to strong opioids. Several national guidelines, including those of Canada, France and Germany adapted this approach for chronic non-cancer pain, recommending treatment initiation with low doses while slowly titrating until adequate levels of analgesia are achieved (Bertin et al., 2026; Busse et al., 2017; Häuser et al., 2014). This conventional escalation approach creates an individual treatment-history that might inadvertently generate negative treatment expectations at treatment onset since initial encounters with the drug are often associated with minimal or no analgesia (Colloca and Benedetti, 2006; Kessner et al., 2013). From a placebo perspective, reversing this approach by starting treatment with a high, effective dose may instead establish strong positive expectations about treatment efficacy, thereby enhancing drug-related placebo effects. The present study therefore tests whether a descending dosing scheme, starting with a high dose, is more effective in enhancing placebo-related analgesia than the conventional ascending dosing scheme that begins with a low dose. To this end, healthy participants were randomly assigned to either a descending or an ascending treatment group and received sham analgesic treatments framed as varying in dose, with groups differing only in the order in which these doses were applied. During the test phase, both groups received an identical treatment dose, enabling a direct comparison of whether treatment-history differentially influenced pain relief between groups.

## Materials and Methods

### Participants

Eighty-eight healthy participants took part in the study. Exclusion criteria included pregnancy, regular drug intake (except contraceptives), psychiatric or neurological disorders, substance abuse, any pain-related disorder and injuries, infections, eczemas or irritations of the skin on the volar forearms. Furthermore, participants were required to abstain from analgesics within 24 hours prior to study participation. Importantly, participants were only invited to the experiment if they had never participated in another placebo-related study within the institute. The study was approved by the Ethics Committee of the Medical Council of Hamburg (PV4817) and conducted according to the Declaration of Helsinki. Written informed consent was obtained from all participants prior to study participation. After study participation, participants received a compensation between 37 and 45€ depending on study duration (approximately 2.5 - 3 hours). Six participants had to be excluded from the sample. Three participants aborted the experiment, one participant had difficulties feeling differences between conditions/temperatures and two participants showed very low ratings across all conditions (see behavioural data analysis). The final sample consisted of 82 participants between 18 and 40 years (mean age 25.8 ± 4.4 years) with 37 male and 45 female participants. This final sample was composed by pooling data from two independent samples to increase statistical power. Sample 1, which was collected in 2019 and comprised 20 participants, served as pilot study for sample 2, which was collected in 2022 and comprised 62 participants. In sample 1, we implemented a within-subject study design were participants experienced both treatment-history sequences (ascending and descending) in counterbalanced order. However, because of carry-over effects, only the first treatment-history sequence was included in the analysis, therefore treating sample 1 as a between-subject comparison. As a consequence, sample 2 was collected using a between-subject design, with participants being randomly assigned to one of the two treatment-history sequences. Experimental differences between the two samples are noted in the methods section where relevant.

### Study design

The aim of the study was to investigate influences of treatment-history on subsequent treatment-related pain relief. Therefore, we developed a heat pain paradigm (Fig. 1A) with different analgesic treatment conditions where a sham analgesic treatment was administered at three different “doses” (low, medium and high) to obtain different levels of pain relief in comparison to a no-treatment condition (control). During the treatment-history phase, two temporal sequences of treatment conditions were applied in a between-subjects design while in the test phase, all participants received the same treatment condition. Before the experiment started, every participant was calibrated to obtain four temperatures that matched 25, 40, 55 and 70 points on a Visual Analogue Scale (VAS). Participants were then randomly assigned to one of the two treatment-history groups. The descending treatment group (N = 40) received a temporal sequence with treatment conditions that decreased in its analgesic efficacy (high, medium, low) while the ascending treatment group (N = 42) received a temporal sequence with treatment conditions that increased in its analgesic efficacy (low, medium, high). Note that groups were named according to the direction of analgesic efficacy, not stimulus intensity; for instance, the descending group's dose decreased over time, implemented through an ascending stimulus intensity which is the reverse of what the group name suggests. As sham analgesic treatment, we introduced transcutaneous electrical nerve stimulation (TENS) that was applied around the thermal stimulation area. Participants were instructed that the frequency of TENS influences the level of pain relief similar to different doses of an analgesic drug with higher frequencies yielding higher pain relief. Hence, for the three treatment conditions, three different TENS frequencies (50, 100, 150 Hz) were supposedly applied to obtain different levels of pain relief while no TENS (0 Hz) was applied during the control condition to let participants experience the pain stimulation without treatment (Fig. 1B). Importantly and unbeknownst to the participants, the TENS stimulation was not applied during the treatment conditions but the temperatures for the low, medium and high treatment conditions were lowered to correspond to 55, 40 and 25 VAS, respectively while the temperature in the control condition corresponded to 70 VAS. The reason for implementing sham treatments with covertly decreased temperatures instead of real analgesic drugs was to overcome the typically large interindividual variability in drug responses and to keep the logistical burden of the experiment within a reasonable range.

**Figure 1.**
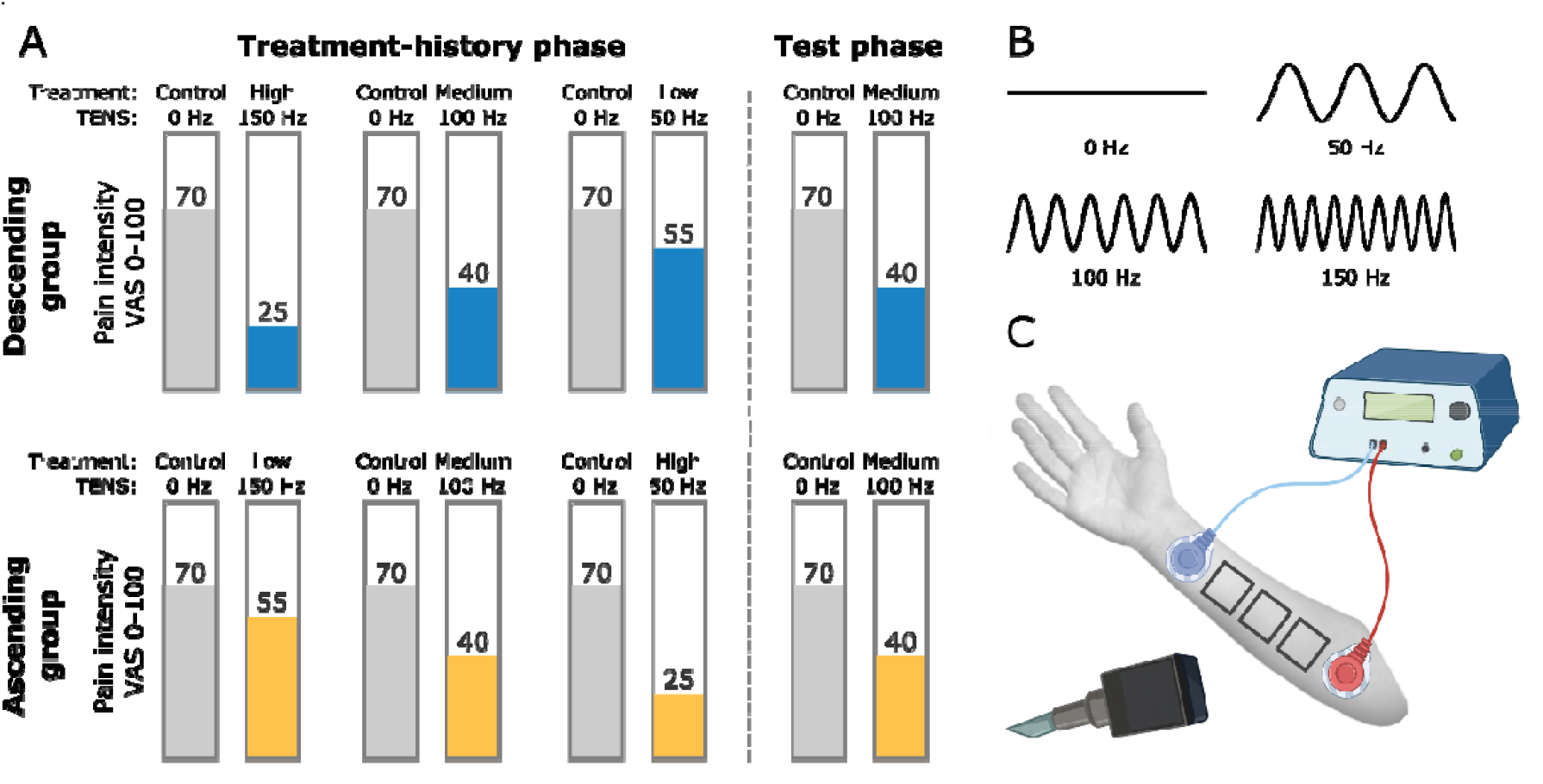
Study design. **(A)** Experimental procedure for both treatment groups with intended pain intensity levels. The descending treatment group (blue) received a descending dosing scheme that started with a high treatment while the ascending treatment group (yellow) received an ascending dosing scheme that started with a low treatment. During the test phase, both groups both groups received the same medium treatment condition. Note that the terms descending and ascending refer to the order of treatment doses, not stimulus intensity, i.e. a high treatment is represented through a low pain intensity level. **(B)** Frequency cues for the control condition (0 Hz) and the three treatment conditions (50, 100, 150 Hz) that were presented in the beginning of every microblock. **(C)** Experimental setup with thermode and TENS stimulation on the volar forearm.

The treatment-history phase consisted of three blocks and every block consisted of two microblocks. The first microblock represented always the control condition and the second microblock one of the three treatment conditions. The control condition preceded every treatment condition to account for temporal effects such as habituation or sensitization which could otherwise confound ratings between treatment groups. In sample 1, every microblock comprised six stimuli while in sample 2, every microblock comprised eight stimuli. In the beginning of every microblock, a cue indicated the TENS frequency that would supposedly be applied during the microblock (Fig. 1B). In microblocks with the control condition, a line representing 0 Hz was used as cue. After the treatment-history phase, all participants underwent the same test phase that comprised a fourth block with a control condition and a subsequent medium treatment condition.

### TENS cover story

The TENS device was a direct current stimulator (Digitimer Constant Current Stimulator, Digitimer) that delivered electrical stimulation to two electrodes that were attached above and below the three marked skin patches on the left volar forearm (Fig. 1C). Participants were informed that TENS is a device used in pain treatment that reduces afferent nociceptive signaling. Participants were further instructed that different levels of pain relief can be obtained by applying different frequencies of the electrical stimulation. To let participants experience the TENS stimulation, a few electrical stimuli were applied throughout the experiment. After heat calibration, participants were told that their TENS detection threshold would be measured. Therefore, one stimulus with 0 µA was applied, another with 50 µA that most participants perceived and again a stimulus with 0 µA. In case that participants did not perceive the second stimulus, the intensity was increased in increments of 10 µA until the participant perceived the electrical stimulation. Moreover, after every skin patch change two electrical stimuli were applied which was introduced as a recalibration of the TENS stimulation. Importantly, participants were informed that the stimulation during the experiment was below their perception threshold, which is why they should not expect to feel the TENS stimulation during the experiment. After the completion of the experiment, participants were informed about the deception of the TENS stimulation. Note that the experimenter changed between sample 1 and 2. Although we tried to keep the TENS-related information as consistent as possible between experimenters, different wording or emphasis might still have occurred between samples.

### Thermal stimuli and trial design

A CHEPS Thermode (27 mm surface diameter, Medoc, Ramat Yishai, Israel) delivered thermal stimuli to participants’ left volar forearm. The baseline temperature was set to 34°C and thermode rise and fall rates were 15°C/s. During the experiment, three different skin patches were stimulated. One skin patch was stimulated during the calibration phase, the second during the conditioning phase and the third during the test phase. The order of skin patch stimulation was counterbalanced between participants. With regard to the trial design, each trial consisted of a jittered anticipation phase (1.5 - 2.5 s) that started with a crosshair changing its colour from white to red followed by a painful stimulus (6 s plateau) where the crosshair remained red. During the rating phase (5 s) a visual analogue scale appeared on the screen and participants assessed the painfulness of the preceding stimulus. The scale consisted of 101 rating steps from 0 to 100. The left anchor of the scale was labelled with ‘no pain’ and the right anchor was labelled with ‘unbearable pain’. During the jittered inter-stimulus interval (11 - 15 s), participants saw a white fixation cross. Stimulus presentation, thermode triggering and response logging were realized with MatlabR2017a and Psychophysics Toolbox (http://psychtoolbox.org). Due to a thermode issue, affected trials were excluded from analyses, which represented 1.9% of all trials.

### Calibration

Prior to the experiment, temperatures were calibrated to infer four different temperatures for every participant that would be rated around 25, 40, 55, and 70 VAS. The calibration procedure consisted of three phases. In the first phase, individual pain thresholds were determined using a probabilistic procedure (Awiszus, 2003). Here, participants indicated whether heat stimuli were ‘painful’ or ‘not painful’. For the second phase, participants rated heat stimuli of different intensities that were estimated based on their pain threshold on a VAS scale from 0 (no pain) to 100 (unbearable pain). In sample 1, participants received 16 heat stimuli between -2 and +2.5 °C around their pain threshold in a pseudorandomized order. In sample 2, an adaptive procedure was used to estimate individual pain windows. Here, four stimuli were applied that increased by 0.5, 1, 2 and 2.5°C based on the pain threshold. These ratings were used to infer twelve temperatures that participants would rate between 10 and 90 VAS and subsequently applied in pseudorandomized order. In both samples, all ratings from the second phase were used to determine four temperatures that matched ratings of 25, 40, 55, and 70 VAS. In the third phase, the four temperatures were applied twice to ensure that participants were well calibrated and noticed differences between the temperatures as this was essential for the experiment. In case that ratings between the four temperatures did not differ sufficiently, temperatures were slightly adapted.

### Questionnaires

Participants completed a set of questionnaires after the experiment. The set included the Beck Depression Inventory (BDI), the State and Trait Anxiety Inventory (STAI), and the Social Desirability Scale (SDS). After the experiment, an exit questionnaire assessed whether participants felt differences between the three treatment conditions. Therefore, participants provided efficacy ratings for every treatment condition between 0 and 100 (handwritten numbers). Participants further indicated whether they would use TENS as analgesic treatment and/or recommend it to a friend with the aim to indirectly assess whether participants believed our TENS-related cover story.

### Behavioral data analysis

Behavioral data were visualized and analyzed in MATLAB (version R2019b; The MathWorks Inc.) and R (version 4.6.1; R Core Team, 2026). Two of the six excluded participants were flagged during inspection of the treatment-history data as their average rating in the control condition was below 35 VAS across all blocks. Similarly, all three treatment conditions were rated below 10 VAS, indicating floor effects. We therefore excluded these participants from all following analyses. Within the final sample, we compared groups and samples with regard to demographical variables in a first step. To this end, linear regression models (lm in R) were computed for continuous variables and logistic regression models (glm (binomial) in R) for categorical variables. In a second step, we analyzed whether groups differed in their pain calibration (temperatures, ratings) using linear mixed models (lmer in R). Group differences in calibrated temperatures were modelled using the four temperatures as dependent variable, intensity level (25, 40, 55, 70 VAS) and group as fixed effects and subject number as random effect. Group differences in ratings (provided at the end of the calibration phase) were modelled using the four averaged ratings as dependent variable, intensity level (25, 40, 55, 70 VAS) and group as fixed effects and subject number as random effect. Similarly, all research questions regarding pain ratings during the experiment were analyzed using linear mixed models (lmer in R). The factor sample was included as a covariate in all models (including models on calibration data) to account for procedural differences between the two pooled samples. Post-hoc comparisons were realized though marginal means (emmeans in R) and subsequent contrast estimations of interest. Post-hoc tests were corrected for multiple comparisons using the Holm method.

The treatment effect in the treatment-history phase was modelled using a linear mixed effects model with trial-wise pain ratings as dependent variable, condition (control, treatment), level (high, medium, low) and group as fixed effects and subject number as random effect.

#### Rating ∼ Condition ^*^ Level ^*^ Group + Sample + (1| Subjects)

Post-hoc tests compared differences between groups within one level between control and treatment ratings. The treatment effect in the test phase was modelled using a linear mixed effects model with trial-wise pain ratings as dependent variable, condition (control, treatment) and group as fixed effects and subject number as random effect.

#### Rating ∼ Condition ^*^ Group + Sample + (1| Subjects)

A post-hoc test was performed to compare group differences between control and treatment ratings. To investigate whether the treatment protocol also had an influence on the control condition, we compared ratings in control microblocks during the treatment-history phase. A linear mixed model was implemented with trial-wise pain ratings as dependent variable, block (1 to 3, temporal order) and group as fixed effects and subject number as random effect. Post-hoc tests were performed to test for baseline differences between groups (block 1) and to test whether control ratings changed compared to baseline between groups (blocks 2 and 3). Finally, for the treatment efficacy ratings provided in the exit questionnaire, a linear mixed model was computed with efficacy ratings as dependent variable, frequency cue (50, 100 150 Hz) and group as fixed effects and subject number as random effect. Moreover, answers to the questions whether participants would use TENS themselves or recommend it to friend were grouped into three categories (yes, no, maybe) and multinomial logistic regression models (multinom in R) were computed to compare groups.

## Results

### Calibration

Groups did not significantly differ in age, gender, handedness, pain thresholds, depressive symptoms, state and trait anxiety, or social desirability (Table 1). Moreover, there were no significant differences between the two samples for these variables.

**Table 1.** Sample characteristics. Group and sample comparisons based on demographical variables, pain thresholds and questionnaire scores. Values represent the group mean ± standard deviation (columns 1 and 2) and statistical values for group comparison (column 3) and sample comparison (column 4).

| | Descending<br>Group (N = 40) | Ascending<br>Group (N = 42) | Group<br>F/$\chi^2$ (p-value) | Sample<br>F/$\chi^2$ (p-value) |
| --- | --- | --- | --- | --- |
| Age (years) | 25.9 ± 4.7 | 25.8 ± 4.1 | 0.02 (0.90) | 0.006 (0.94) |
| Gender (male/female) | 17/23 | 20/22 | 0.32 (0.57) | 0.03 (0.87) |
| Handedness (left/right) | 3/37 | 7/35 | 1.76 (0.19) | 0.17 (0.68) |
| Pain Threshold ($^{\circ}$C) | 43.9 ± 0.7 | 43.6 ± 1.0 | 2.78 (0.10) | 2.58 (0.11) |
| BDI | 5.9 ± 4.9 | 7.4 ± 6.0 | 1.70 (0.20) | 0.07 (0.79) |
| STAI-State | 35.0 ± 8.0 | 34.6 ± 8.3 | 0.02 (0.89) | 0.20 (0.66) |
| STAI-Trait | 37.5 ± 8.4 | 36.6 ± 10.6 | 0.10 (0.75) | 0.05 (0.83) |
| SDS | 13.4 ± 3.8 | 12.9 ± 4.2 | 0.42 (0.52) | 1.68 (0.20) |

Linear mixed models investigating whether there were group differences in calibrated temperatures revealed no significant difference between groups (F[1,79] = 0.78, p = 0.38) and no significant group by intensity level interaction (F[3,240] = 0.09, p = 0.96, Fig. 2A). Importantly, ratings provided for the four intensity levels during the calibration procedure indicate that on average participants rated the stimuli close to the intended pain intensity levels (Fig. 2B). A linear mixed model testing for group differences in these ratings did not show any difference between groups (F[1,79.08] = 0.65, p = 0.42) nor any significant interaction between group and intensity level (F[3,239.19] = 0.07, p = 0.98). With regard to samples, there was a significant difference between samples in calibrated temperatures (F[1,79] = 7.49, p = 0.0077, Fig. S1A). On average, temperatures were 0.7°C lower in sample 1 compared to sample 2, likely due to the different thermodes used to stimulate the two samples. Importantly, there was no significant difference in ratings between the two samples (F[1,79] = 0.001, p = 0.97, Fig. S1B), indicating that both calibration procedures were successful in reaching target VAS ratings albeit differences in absolute temperatures.

**Figure 2.**
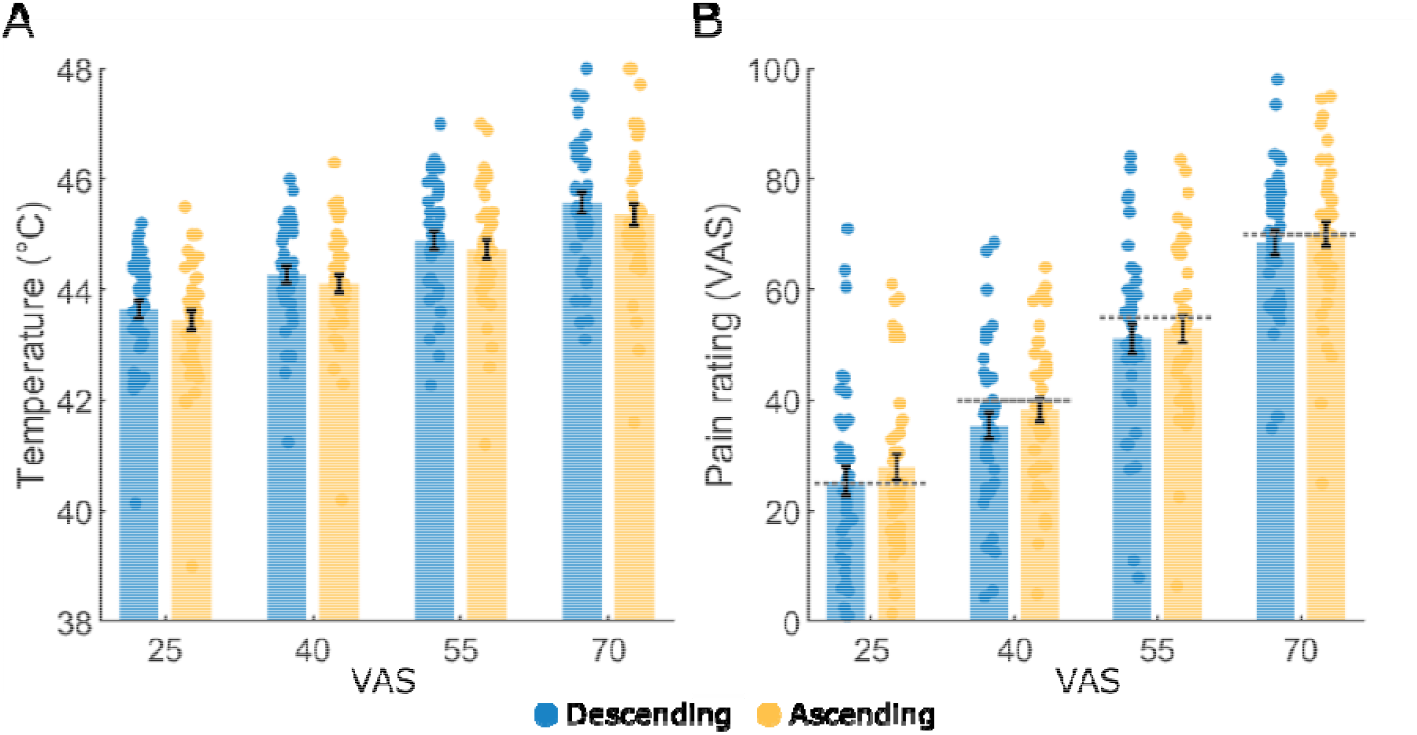
Calibration results. **(A)** Mean calibrated temperatures for every intensity and group. **(B)** Mean VAS ratings, provided after the calibration phase and before the start of the experiment, for every intensity level and group. The dashed lines indicate the intended intensity ratings. Error bars represent the standard error of the mean and data points reflect individual participants.

### Experiment

For visualization purposes, mean pain ratings for every microblock and treatment group are shown in temporal order in Fig. 3A. Treatment ratings during the treatment-history phase depicting individual trajectories show that most participants clearly perceived the descending/ascending treatment order (Fig. S2). To test our main hypothesis, namely whether a descending dosing scheme increases treatment-related pain relief compared to an ascending dosing scheme, we examined group differences in pain relief between the control and treatment condition during the test phase. This analysis revealed no significant main effect of group (F[1,78.69] = 0.46, p = 0.50) or sample (F[1,80.87] = 0.61, p = 0.44). However, there was a significant group by condition interaction (F[1,1135.8] = 52.60, p < 0.001). A post-hoc comparison confirmed that this effect resulted from the stronger pain relief in the descending group compared to the ascending group (estimate±SE: 12.1 ± 1.66, t_1136_ = 7.25, p < 0.001, Cohen’s d = 0.83, Fig. 3B), indicating that a descending dosing scheme enhances treatment-related analgesia. Based on this result, we next asked whether these group differences in treatment-related pain relief were already present during the treatment-history phase. We therefore tested whether pain relief differed between groups across treatment levels (Fig. 3C). This analysis again revealed no significant main effect of group (F[1,78.8] = 0.27, p = 0.60) or sample (F[1,79.7] = 0.09, p = 0.76). However, significant two way interactions emerged between group and condition (F[1,3530.8] = 29.25, p < 0.001) and group and level (F[1,3530.8] = 8.72, p < 0.001). Post-hoc comparisons testing for group differences in pain relief at each treatment level showed that the descending group already exhibited stronger pain relief during the treatment-history phase than the ascending group across all three treatment levels: low treatment (estimate±SE: 4.45 ± 1.59, t_3531_ = 2.80, p = 0.01, Cohen’s d = 0.32), medium treatment (estimate±SE: 6.85 ± 1.60, t_3531_ = 4.29, p < 0.001, Cohen’s d = 0.50), high treatment (estimate±SE: 3.64 ± 1.6, t_3531_ = 2.28, p = 0.023, Cohen’s d = 0.26).

**Figure 3.**
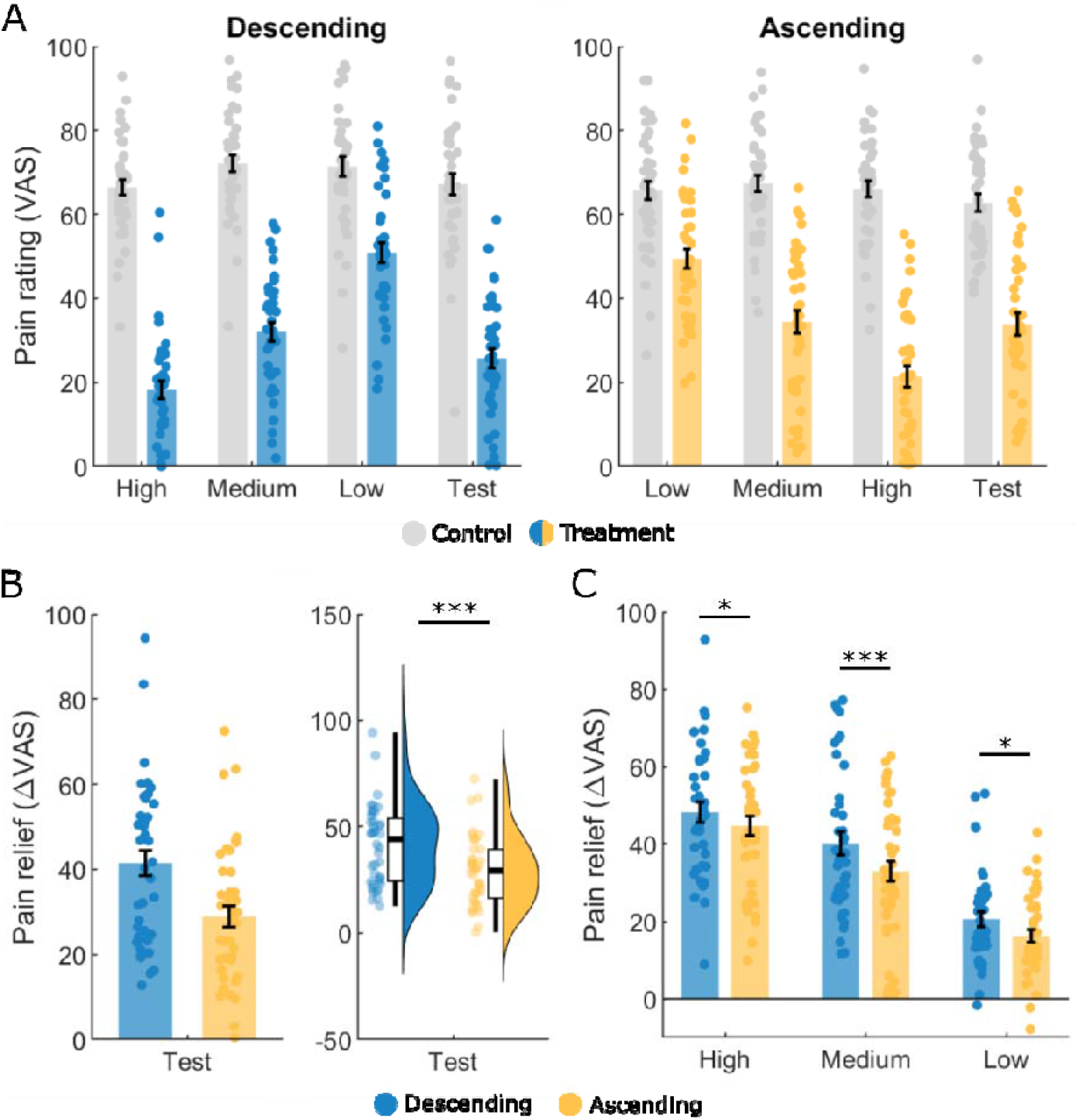
Treatment-history results. **(A)** Mean pain ratings in temporal order for every microblock and group (left panel: descending group; right panel: ascending group). **(B)** Mean pain relief (control-treatment) in the test phase for the medium treatment condition in both groups. Left panel: bar plot; right panel; box plot with density distribution. **(C)** Mean pain relief in the treatment-history phase, expressed as difference between control and treatment microblock, for every treatment condition and group. Error bars represent the standard error of the mean and data points reflect individual participants. Asterisks indicate significant differences (^*^p < .05, ^***^p < .001).

To further assess whether these group differences during the treatment-history phase emerged primarily in the treatment conditions, or whether the control condition was also influenced by the dosing scheme, we exploratorily analysed group differences in control ratings across the three treatment-history blocks. This analysis revealed a significant block by group interaction (F[2,1732.76] = 5.88, p = 0.0029, Fig. 4A). Post-hoc comparisons showed no significant difference between groups in block 1 (estimate±SE: 1.02 ± 2.62, t_94.7_ = 0.39, p = 0.70, Cohen’s d = 0.09), indicating that groups did not differ at baseline. However, significant group differences emerged in block 2 (estimate±SE: 3.72 ± 1.35, t_1733.1_ = 2.77, p = 0.011, Cohen’s d = 0.32) and block 3 (estimate±SE: 4.18 ± 1.33, t_1733_ = 3.13, p < 0.0053, Cohen’s d = 0.36), relative to block 1, suggesting that following the first treatment experience, the descending group rated the control condition slightly higher than the ascending group.

**Figure 4.**
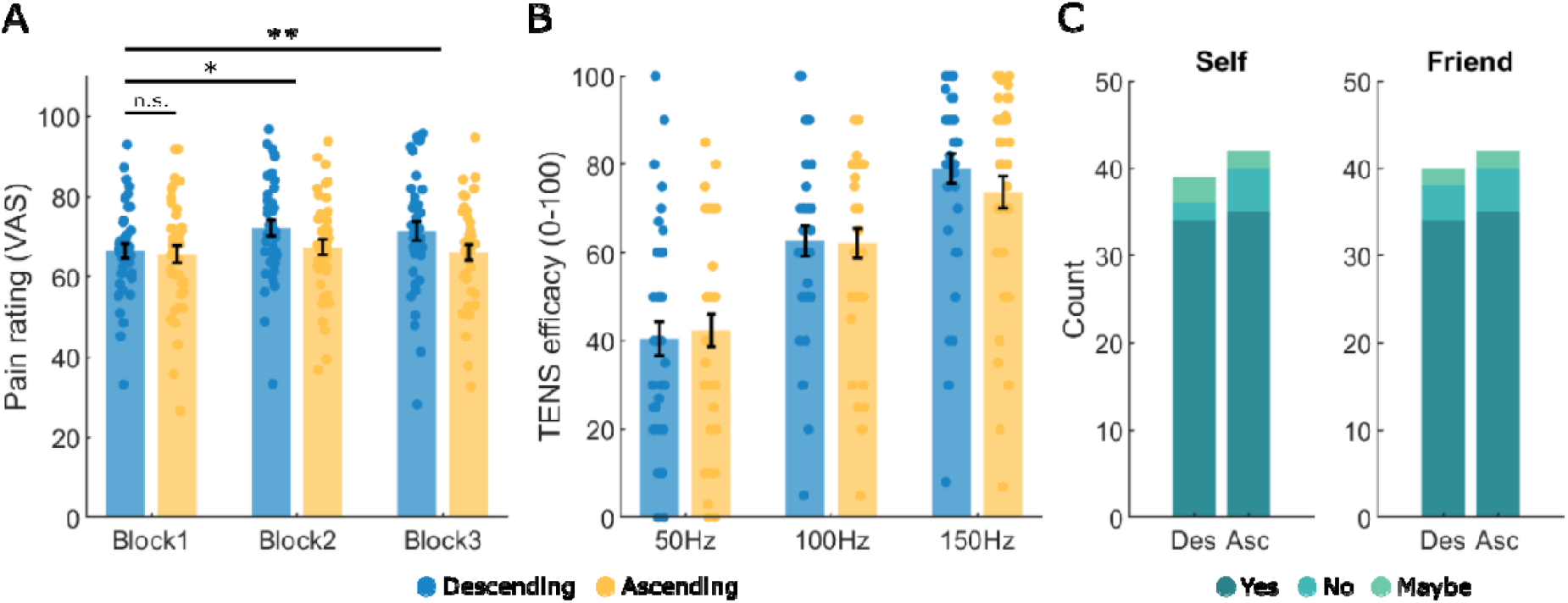
Additional results. **(A)** Mean pain ratings in the control condition for every treatment-history block and group. **(B)** Handwritten numeric mean TENS efficacy ratings for every treatment condition and group, provided after the experiment. **(C)** Distribution of answers (yes, no, maybe) per group to the question whether participants would use TENS for themselves (left panel) and/or recommend to a friend (right panel). Error bars represent the standard error of the mean and data points reflect individual participants. Asterisks indicate significant differences (^*^p < .05, ^**^p < .01, n.s.: not significant).

### Additional analyses

After the experiment, participants rated the effectiveness of the three TENS frequencies on a scale from 0 (not effective) to 100 (very effective). These ratings show a clear distinction between the three frequencies (F(2,158) = 123.1, p < 0.001, Fig. 4B, Fig. S3) indicating that participants consciously experienced differences in pain relief across treatment conditions. However, there was no significant difference between groups (F(1,78) = 0.09, p = 0.77) or samples (F(1,78) = 0.15, p = 0.70), nor a significant group by cue interaction (F(2,158) = 1.38, p = 0.26). To assess whether participants believed our TENS stimulation cover story, we asked them whether they would potentially use TENS as analgesic treatment for themselves and/or whether they would recommend it to a friend. This analysis revealed that 79% of participants (N = 65) would both use TENS as possible analgesic treatment and recommend it to a friend. Six participants would either use TENS for themselves or recommend it to a friend while only four participants would neither use it as treatment for themselves nor recommend it to a friend. Importantly, these answers did not differ between groups (self: χ^2^ = 1.57, p = 0.46; friend: χ^2^ = 0.10, p = 0.95) or samples (self: χ^2^ = 3.16, p = 0.21; friend: χ^2^ = 0.05, p = 0.98, Fig. 4C).

## Discussion

This study tested whether the direction of an analgesic dosing scheme influences the strength of expectation-induced pain relief. Contrary to current clinical guidelines that recommend initiating analgesic treatment with a low dose and titrating upward, we found that a descending dosing scheme, starting with a strong, effective dose, produced significantly greater pain relief during a subsequent identical test treatment than the conventional ascending scheme. Notably, this group difference emerged already during the treatment-history phase and was present in all three treatment levels. Together, these findings suggest that the order in which analgesic treatment efficacy is experienced can shape future treatment outcomes.

### Treatment-history effects

Our main results showed that the order in which analgesic efficacy, implemented through different supposedly analgesic doses, can substantially influence the perceived treatment efficacy. Participants who started with a high treatment, with efficacy decreasing over time, reported stronger pain relief in response to a medium test treatment than participants who started with a low treatment, with efficacy increasing over time. These findings complement previous research on treatment-history effects in the placebo field that has predominantly focused on negative treatment experiences, establishing that one negative or ineffective analgesic treatment experience negatively affects subsequent treatment outcomes (Colloca and Benedetti, 2006; Kessner et al., 2014, 2013) that even generalize to different analgesic treatment routes (Zunhammer et al., 2017). Moreover, negative treatment experiences have been shown to impair future treatment success in depression (Ellis et al., 2014; Hunter et al., 2015). These findings show that negative treatment experiences generate negative expectations that are transferred to novel treatment situations and create nocebo-like effects that interfere with treatment success. The ascending treatment group's results can be interpreted in light of negative expectations; even though participants were told they would receive a low-dose treatment, their first analgesic experience provided only minimal relief compared to the control condition. This first negative experience might than have generalized to the other treatment conditions, resulting in reduced pain relief. This interpretation is further supported by an animal study that used a similar ascending dosing scheme in a pharmacological conditioning experiment and did not observe conditioned analgesia after substituting the drug with saline (Boorman and Keay, 2021b). Conversely, the descending treatment group’s first encounter with the analgesic treatment was positive, which might have created strong beliefs about the analgesic efficacy of the treatment that generalized to subsequent treatment conditions. This account speaks for a pure expectation mechanism as the very first encounter sets all expectations that influence all subsequent experiences. However, placebo-like effects in the medical context are usually a combination of expectations and learning so it is plausible to assume that the treatment-history phase represented some kind of conditioning, initiating learning that involves updating of expectations after gathering new information. Another interesting finding was that the control condition was rated as slightly more painful in the descending group than in the ascending group, with this difference emerging after the first treatment experience. This indicates that expectations generated during the first treatment experience also influenced the perception of the no-treatment condition. One possible explanation is a contrast effect, a well-known phenomenon in perceptual judgment (Fritsche et al., 2017; Helson, 1964) whereby a stimulus is perceived as more intense when it follows a stimulus of lower intensity, and less intense when it follows one of higher intensity.

### Clinical implications

These findings raise the possibility that conventional escalation approaches starting treatment with low doses that were primarily designed around safety and tolerability, may carry an unintended cost. By pairing first treatment experiences with minimal to no analgesic efficacy, these conventional approaches may inadvertently generate negative treatment expectations that attenuate the placebo-related component of subsequent treatment responses. Thus, initiating treatment with a stronger, clearly effective dose, could enhance the overall analgesic response to later treatments. Such an approach could, in principle, reduce the overall quantity of analgesic drugs required for pain control, since sufficiently strong positive expectations at treatment onset would allow lower subsequent doses to achieve comparable pain relief. Similar strategies to harness placebo effects for dose reductions in long-term analgesic treatments propose dose-extension paradigms (Colloca et al., 2016; Doering and Rief, 2012) where classical conditioning is implemented through the pairing of the drug with a neutral cue. In the extension phase, the drug is either substituted with a placebo or its dose is reduced, in the hope that the drug effect persists at full efficacy. One striking example of such an approach used an partial-reinforcement schedule with alternating drug and placebo administrations in the extension phase, showing that elevated immune responses persisted even on placebo days (Albring et al., 2014). This has also been demonstrated in patients, with partial reinforcement shown to reduce immunosuppressant dosing in renal transplant recipients while preserving treatment efficacy (Kirchhof et al., 2018). Similar approaches have been investigated in ADHD and psoriasis treatments (Ader et al., 2010; Sandler et al., 2010). In the pain domain, open-label placebos have been used as a conditioned cue paired with opioid administration after surgery. In the two weeks following hospital discharge, patients in the open-label placebo group showed 30% lower opioid consumption than standard care alone (Flowers et al., 2021), suggesting that even without a predefined dose-extension schedule, patients either reduced their opioid dose while taking the placebo or periodically substituted the drug with the placebo. The present findings add a novel dimension to these dose-extension strategies: rather than only maintaining efficacy at a reduced dose after conditioning with a fixed dose, our results suggest that the order of dose exposure itself may be an additional target for enhancing drug-related placebo effects.

### Sample differences

In the present study, we pooled data from two samples to increase statistical power. However, there were minor procedural differences between the two samples. First, the number of trials per microblock differed between the samples (6 vs. 8), which might have impacted learning and expectation generation. Second, the experimenters differed between samples which again might have differentially impacted expectation generation during the cover story instruction. Finally, the calibration procedure differed between samples which in theory could have biased ratings between intensity levels. Intensity levels only differed by 15 VAS points and for the treatment-history phase it was crucial to experience differences between intensity levels/treatment conditions in order to build expectations as intended by the experimental manipulation. Hence, differences in the calibration procedure might have resulted in differing discriminability between the calibrated temperatures, potentially impacting expectations and experiences during the treatment-history. However, we did not observe any significant difference between samples in all our analyses concerning the experimental manipulation, suggesting that the procedural differences did not meaningfully impact expectations and experiences during the experiment. However, we found a statistically significant difference in calibrated temperatures between the two samples, likely due to the different thermodes used to stimulate each sample. Importantly, this difference did not translate into differing pain ratings since average ratings were comparable across intensity levels in both samples, confirming that the calibration successfully achieved its purpose of identifying temperatures that participants would reliably rate at each intended pain intensity level.

### Exit questionnaire

We included an exit questionnaire at the end of the experiment to assess whether participants consciously perceived differences between treatment levels and whether they believed our cover story that TENS is capable of inducing analgesia in the context of heat pain. Most participants consciously experienced that analgesia increased with the supposedly increasing TENS frequency, indicating that our treatment-history phase successfully created the experience of either ascending or descending treatment efficacy. However, there were no group differences in these ratings, which suggests that analgesic differences during the test phase did not transfer to these post-experimental ratings, likely due to the fact that most participants provided (handwritten) numbers in increments of ten. We included two questions in this questionnaire to indirectly assess whether participants believed our cover story. Those questions asked whether participants would use TENS as analgesic treatment themselves and/or recommend it to a friend based on the experience during the experiment. As it is difficult to directly measure whether participants believe the cover story in deception experiments, we opted for these indirect questions as we assumed that participants who would indicate to use it themselves and/or recommend to a friend believed our cover story to a great extent. The majority of participants answered at least one of these questions with “yes,” suggesting that most believed our cover story. For the small number who answered both questions with “no,” however, it remains unclear whether this reflects disbelief in the cover story or simply other reasons for declining TENS, such as an aversion to electrical stimulation or reluctance to purchase or carry such a device. In summary, these data show that most participants believed our deception and believed that they had received differing analgesic doses during the experiment.

### Limitations

Several limitations should be considered when interpreting these findings. First, our paradigm used a sham treatment (TENS) with covertly manipulated stimulus intensities rather than an active pharmacological agent, meaning our findings speak to the psychological mechanism of treatment-history effects but cannot directly establish whether the same pattern would hold with real analgesic drugs, where pharmacodynamic factors (e.g., actual tolerance, sensitization) could interact with placebo-related effects. Second, our sample consisted of healthy participants undergoing experimentally induced acute pain, which may not generalize to clinical populations with chronic pain, altered pain processing, or pre-existing treatment expectations shaped by prior clinical experience. Finally, our design did not include a control group unexposed to any treatment-history manipulation, nor a baseline condition indicating the “true,” unbiased pain relief expected during a medium treatment. This means we can only conclude that one treatment-history scheme is preferable to the other, not determine the direction of the underlying effect: for instance, both groups might have shown a genuine placebo effect of differing strength, or one group might have shown a placebo effect and the other a nocebo effect. Future studies could disentangle these possibilities by including such a control group or baseline condition.

## Conclusion

Our findings suggest that the order in which analgesic treatment efficacy is experienced can shape the magnitude of expectation-induced pain relief, with a descending, high-to-low dosing scheme producing stronger treatment-related analgesia than the conventional ascending approach. These results suggest that treatment-history itself may be a suitable target for enhancing analgesic treatment success which, if validated in clinical context, could contribute to reducing cumulative analgesic and opioid exposure without compromising pain control.

## Acknowledgements

The authors declare that they have no competing interests. All data needed to evaluate the conclusions in the paper are present in the paper and Supplementary Materials. AT and MM were supported by the German Research Foundation (TI 1110/1-1).

