## Supplementary Figures 1-3 for "Starting strong: A descending analgesic dosing scheme enhances pain relief compared to a conventional ascending dosing scheme"

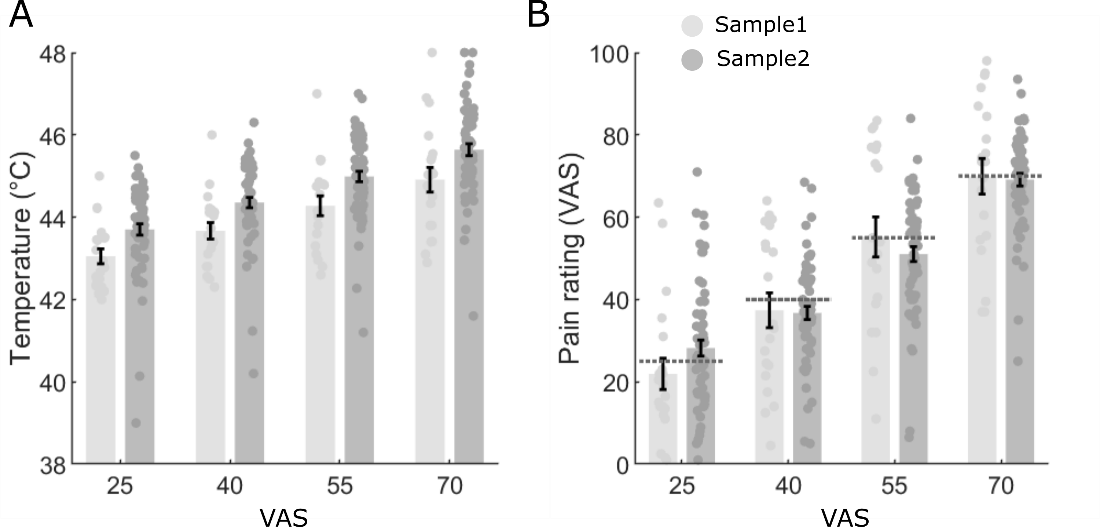

***Figure S1. Sample comparison calibration. (A)*** *Calibrated temperatures for all four pain intensity levels and sample.* ***(B)*** *Mean* *VAS ratings, provided after the calibration phase and before the start of the experiment, for every intensity level and sample. Dashed lines indicate the intended pain rating. Error bars represent the standard error of the mean and data points reflect individual participants.*

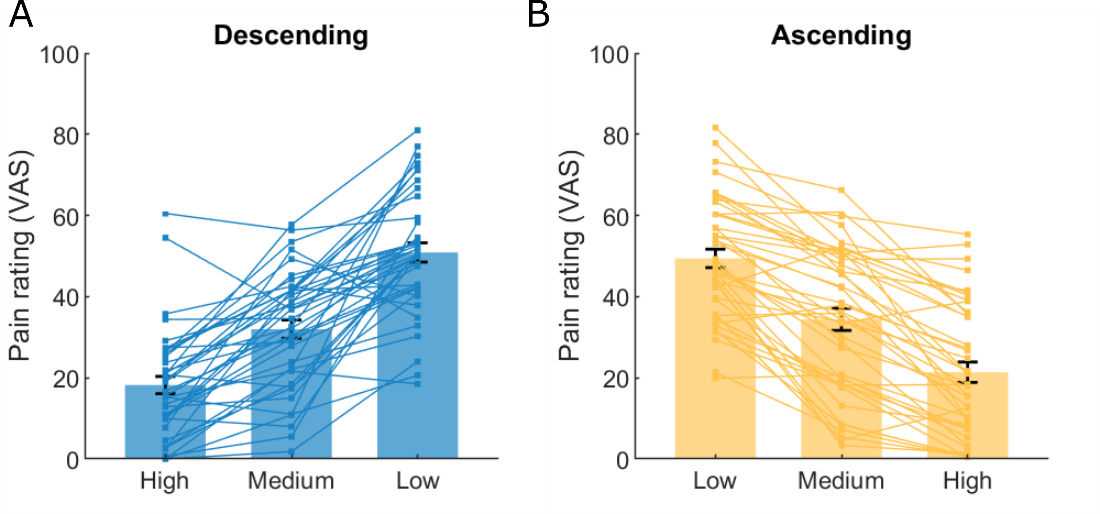

***Figure S2. Individual treatment trajectories during the treatment-history phase. (A)*** *Mean* *VAS ratings in the descending treatment group for all three treatment conditions during the treatment-history phase with individual trajectories.* ***(B)*** *Mean* *VAS ratings in the ascending treatment group for all three treatment conditions during the treatment-history phase with individual trajectories. Error bars represent the standard error of the mean and individual lines reflect individual participants.*

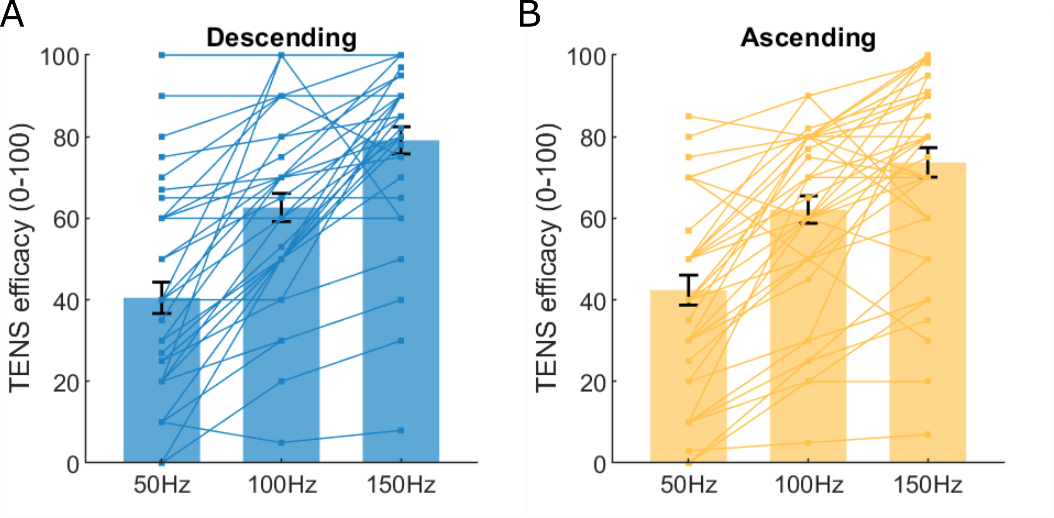

***Figure S3. TENS efficacy ratings with participant trajectories. (A)*** *TENS efficacy ratings in the descending treatment group for all three treatment cues showing individual trajectories.* ***(B)*** *TENS efficacy ratings in the ascending treatment group for all three treatment cues showing individual trajectories. Error bars represent the standard error of the mean and lines represent individual participants.*
